# Structured illumination microscopy combined with machine learning enables the high throughput analysis and classification of virus structure

**DOI:** 10.1101/266551

**Authors:** Romain F. Laine, Gemma Goodfellow, Laurence J. Young, Jon Travers, Danielle Carroll, Oliver Dibben, Helen Bright, Clemens F. Kaminski

## Abstract

Optical super-resolution microscopy techniques enable high molecular specificity with high spatial resolution and constitute a set of powerful tools in the investigation of the structure of supramolecular assemblies such as viruses. Here, we report on a new methodology which combines Structured Illumination Microscopy (SIM) with machine learning algorithms to image and classify the structure of large populations of biopharmaceutical viruses with high resolution. The method offers information on virus morphology that can ultimately be linked with functional performance. We demonstrate the approach on viruses produced for oncolytic viriotherapy (Newcastle Disease Virus) and vaccine development (Influenza). This unique tool enables the rapid assessment of the quality of viral production with high throughput obviating the need for traditional batch testing methods which are complex and time consuming. We show that our method also works on non-purified samples from pooled harvest fluids directly from the production line.

## Introduction

The potential of super-resolution microscopy (SRM) to unravel details of the structure and replication of viruses was recognised early on in the development of the methodology (Betzig et al., 2006; Müller and Heilemann, 2013). Since then, SRM has been used to provide unprecedented insights into viral protein architecture (Laine et al., 2015; Zhang et al., 2015; Gray et al., 2016; Albecka et al., 2016). Previous work has focused on those SRM techniques that achieve the highest theoretical resolution, such as Stimulated Emission Depletion (STED) (Hell and Wichmann, 1994) and Single Molecule Localisation Microscopy (SMLM) (Rust et al., 2006; Heilemann et al., 2008). Whilst offering high fidelity data, the downside is the associated long acquisition time required by these methods, limiting their application to the imaging of static samples at low throughput. A much faster technique, although inferior in spatial resolution, is Structured Illumination Microscopy (SIM) (Gustafsson, 2000; Heintzmann and Cremer, 1999) and this has been applied to study large viruses such as the prototypic poxvirus (Gray et al., 2016; Horsington et al., 2012). In addition to understanding the structure of viruses, there is also need to identify and analyse classes of structures within large viral populations, especially in the biotechnology industry where virus quality is often compromised by large scale production operations and the virus product is often characterised by significant morphological heterogeneities. In particular, campaigns of influenza immunization rely heavily on the timely and efficient production of specific virus strains. Similarly, a deeper understanding of the structural heterogeneity of oncolytic viruses such as Newcastle Disease Virus (NDV) (Ganar et al., 2014; Lichty et al., 2014) would enable optimization of the production processes and in turn improve the development of viriotherapy. However, quantifying and understanding this structural heterogeneity and relating it to virus efficacy requires the imaging of large numbers of viruses at sufficient spatial resolution to reveal characteristic morphological details. Typically, this is achieved by extracting batches from the production process, with elaborate subsequent purification and preparation steps before characterisation by electron microscopy (EM). The method is too slow, low in throughput, and unspecific (for example to discern the presence of particular proteins in the virus envelope) to be of practical use during production operations.

Here, we demonstrate that rapid high resolution imaging with Total Internal Reflection SIM fluorescence microscopy (TIRF-SIM) (Shao et al., 2011; Kner et al., 2009; Young et al., 2016), combined with a machine learning (ML) approach to analyse and classify structures in virus batches offer a great opportunity to circumvent these problems. We present MiLeSIM (Machine Learning Structured Illumination Microscopy) as an efficient combination ofSRM, ML-based classification (Van Valen et al., 2016; Sommer et al., 2011) and advanced image analysis for the quantification of morphological heterogeneities in large virus populations. We use ML algorithms to perform a classification of super-resolved images of a heterogeneous virus population into particle classes with distinct and characteristic structural features (e.g. spherical, filamentous). The classified subpopulations are then further analysed through image analysis pipelines that are specifically adapted for each structural class. We and others have shown that appropriate model fitting can lead to precision in structural parameters beyond the resolution of the images used (Laine et al., 2015; Manetsberger et al., 2015). The method combines speed and specificity and allows an in-depth exploration of large virus populations that is unachievable by electron microscopy (EM). The method has potential in the industrial production of viruses e.g. for oncolytic viriotherapy and vaccine development.

First, we compare TIRF-SIM with alternative imaging modalities and show that it is the method of choice to investigate virus structure at high-throughput (~500 virus particles/s) with a spatial resolution reaching ~90 nm. The large datasets obtained with TIRF-SIM were then fed into an ML algorithm for the automated classification of Newcastle Disease Virus (NDV) and live attenuated influenza virus (LAIV) vaccines, enabling further shape-specific quantitative analyses for a structural description of viral subpopulations. The purpose of our study is to validate the MiLeSIM approach as a powerful analysis tool for biotechnological processes involving virus production both in industry and in the research laboratory.

## Results

### TIRF-SIM offers an optimal combination of throughput and resolution for the imaging of virus structure

First, we explored and compared three common SRM modalities for the structural investigation of purified NDV virus, namely direct stochastic optical reconstruction microscopy, a’STORM, stimulated emission depletion microscopy, STED and TIRF-SIM. Figure 1 shows representative images obtained with all 3 techniques. For comparison, a conventional (non-super-resolved) TIRF wide-field image is also shown. A comparison of performance parameters (resolution and imaging speed) for the different methods is presented in ***Figure Supplement 1***(a). It is clear, that improving resolution beyond the ~90 nm offered by TIRF-SIM comes at a significant cost in acquisition times and throughput.

**Figure 1.**
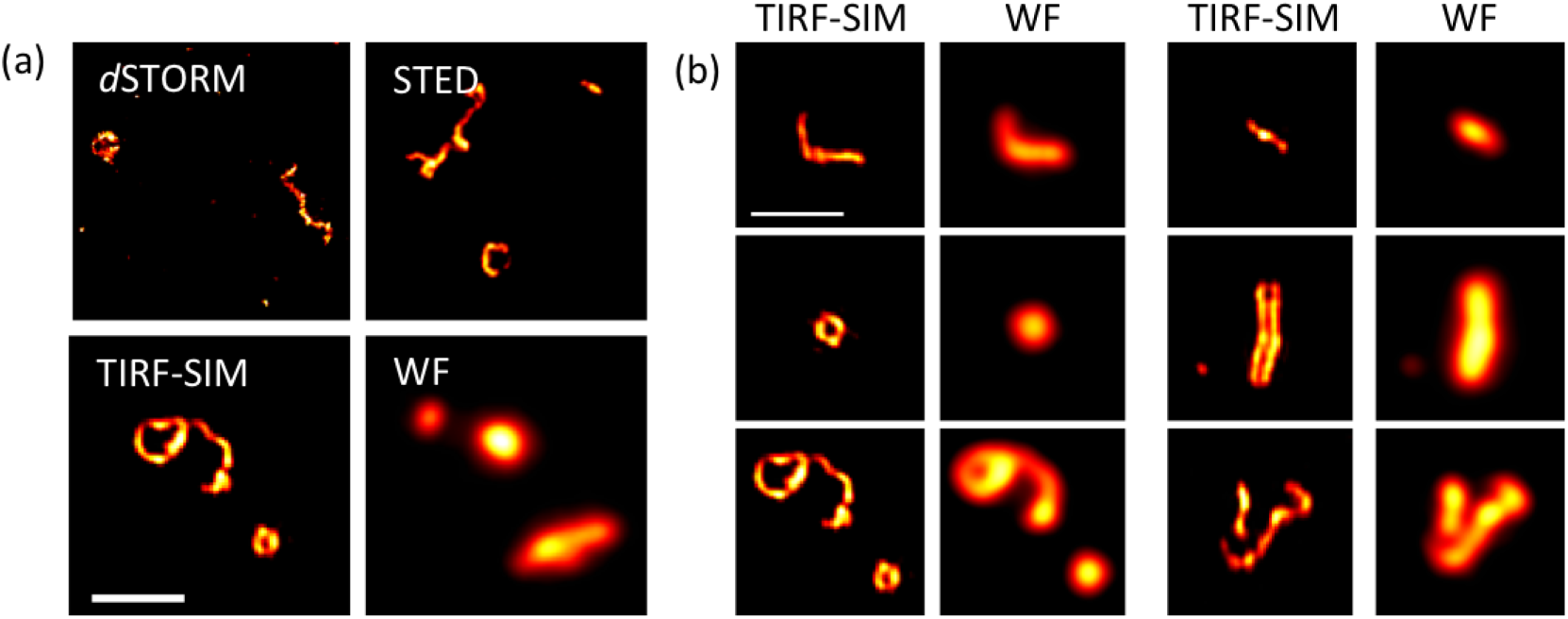
Super-resolution microscopy (SRM) for the study of NDV virus structure. (a) Representative images of purified NDV viruses with different imaging modalities. (b) Representative images of NDV virus population imaged with TIRF-SIM and their corresponding TIRF wide-field image. WF: wide-field TIRF microscopy. Scale bar: 1*μm*.

Furthermore, although *d*STORM and STED offer theoretically higher resolution than SIM, the images obtained with these methods appear not to be of an obviously higher quality than those recorded with TIRF-SIM, and do not reveal additional structural details that are not also resolved by TIRF-SIM images. We conclude that TIRF-SIM is perfectly suited to the present application, providing clear structural details to discern filamentous, spherical and rod-like structures in large NDV populations. Typical shapes discerned within the NDV samples by the method are shown in Figure 1(b).

### Workflow of MiLeSIM

The images obtained with TIRF-SIM show a number of stereotypical virus structures in NDV samples, indicating a large morphological diversity in the virus populations that may stem from variability occurring during viral replication or at the purification stage. Understanding the origins and consequences of such heterogeneity informs not only on the life cycle of the virus but can also provide essential insights into the virus production process to manufacturers of virus-based therapeutics. An automated classification of virus shapes would enable the quantification of virus heterogeneity and permit further analysis of each individual class independently. The workflow to achieve these goals is shown in Figure 2. Individual virus particles are first identified by automated segmentation and then fed to the ML routine for classification. We used a supervised ML algorithm (here a random forest algorithm (Breiman, 2001)) to ensure the robustness of the method and for ease of implementation. We identified 6 major structural classes in the NDV samples which we divide into long and short filamentous, small and large spherical, rod-like and unknown structures. The unknown class is made of clumps of viral material with no consistent and identifiable shapes.

**Figure 2.**
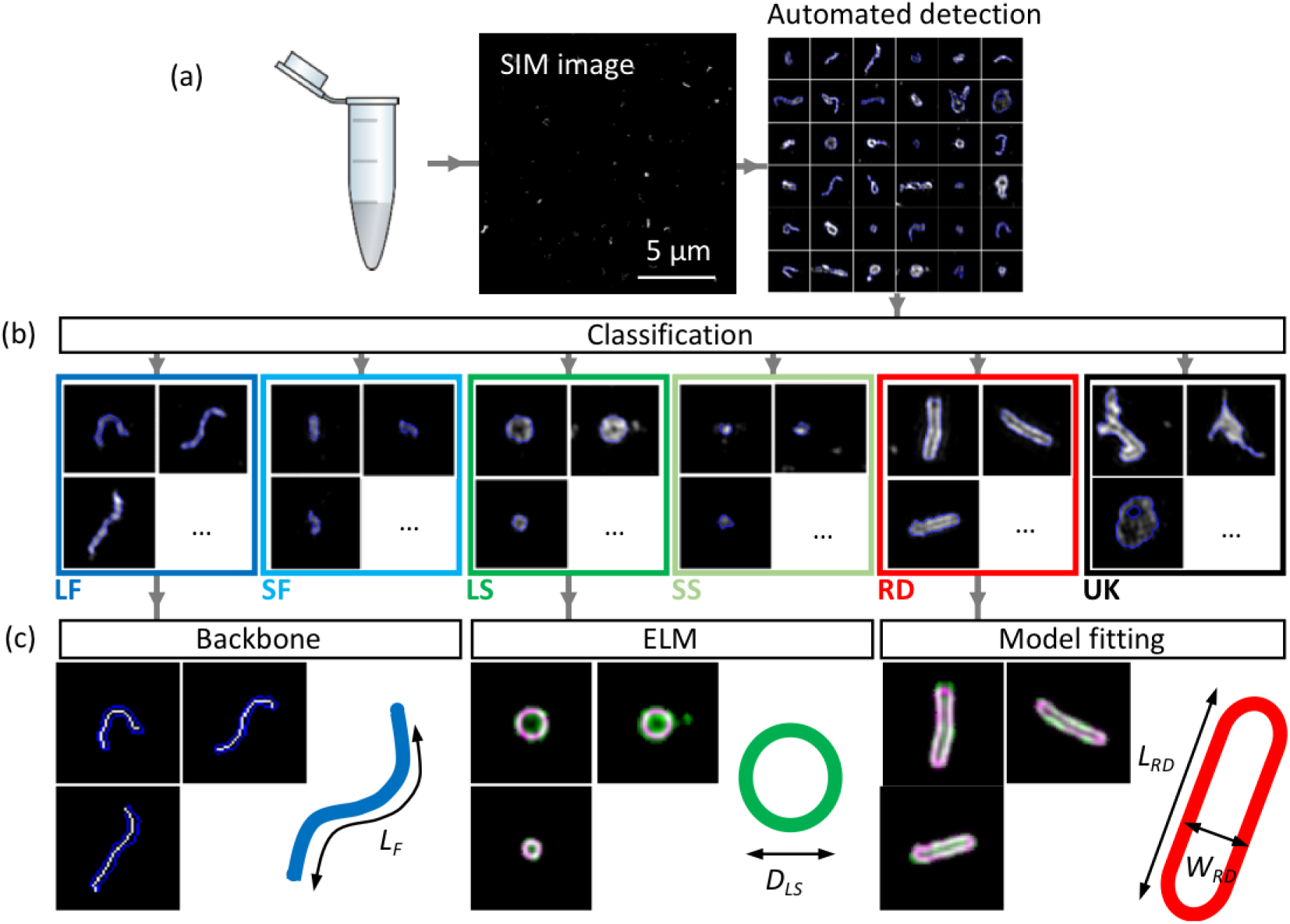
Workflow of automated detection, classification and analysis of NDV viral particles. (a) SIM image and segmentation of individual particles. (b) Classification using ML. (c) The classified single-virus images can be further analysed with a set of class-specific tools. For the backbone analysis the mask and backbone are showed in blue and white respectively. For the model-fitting approach (spherical and rod-like), the data and model are showed in green and magenta respectively. LF: long filamentous, SF: short filamentous, LS: large spherical, SS: small spherical, RD: rod-shape, UK: unknown. *L_F_*, *D_LS_*, *L_RD_* and *W_RD_* represents the length of the filamentous particles, the diameter of the large spherical, and the length of the rod-shaped particles and the width of the rod-shaped particles respectively. Images of individual particles cover a field of view of 1.6 × 1.6 *μm*^2^.

The filamentous particles (long and short) class is further analysed by automatic extraction of the linear backbone of structures and measurement of their length. The width of these filamentous structures appeared to be limited by the resolution of the imaging technique (~90 nm for TIRF-SIM, see ***Figure Supplement 1***, but also observed in higher resolution approaches such as *d*STORM) and therefore, we considered the filamentous class as 1D structures. The large spherical structures were analysed by ellipsoid localization microscopy (ELM) analysis (Manetsberger et al., 2015). The latter fits a shape model to imaging data to permit the extraction of structural parameters with precision higher than the inherent resolution of the imaging method (Laine et al., 2015; Manetsberger et al., 2015). We observed no significant ellipticity in the virus particles and fitted spherical shapes to extract the radius of the particles. A similar model-based fitting approach was used to fit rod-like viral particles and to obtain length and width parameters for this structural class (see materials and methods section and ***Figure Supplement 2*** for details).

### Classification of virus structures using supervised machine learning algorithms

The structural classification was performed using a supervised ML algorithm which allows for rapid and automated classification of large datasets. The choice of algorithm and the set of features (also often called predictors) extracted for each identified particle were optimised to maximise the overall predictive power based on the training dataset (comprising of 370 manually annotated particles). Figure 3 describes the list of chosen individual features (selected from basic shapes features, Hu’s image moments (Hu, 1962), features obtained from the pre-trained convolutional neural network (CNN) AlexNet (Krizhevsky et al., 2012) and from Speeded Up Robust Features, SURF (Bay et al., 2008)) as well as the predictive power associated with the resulting classifier.

**Figure 3.**
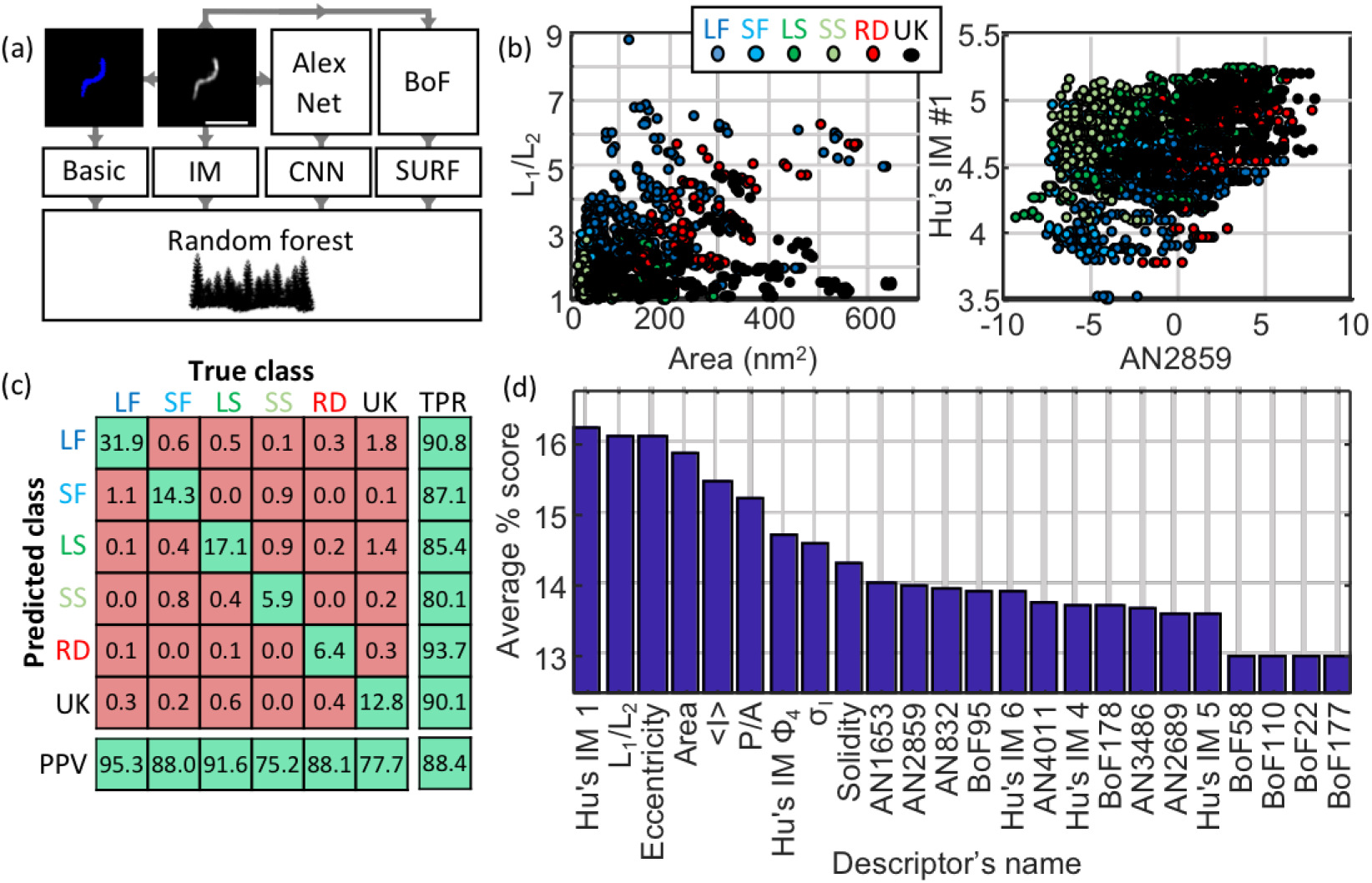
Machine learning-based classification. (a) Building the list of predictors from basic features, image moments, convolutional neural network (CNN) features and SURF bag of features (BoF). (b) 2D scatter plot of pairs of predictors showing the different predictive power of combination of predictors. (c) Confusion matrix obtained from the random forest showing the high true positive rate (TPR) and positive predictive values (PPV) of the classification. All numbers shown here are in percentage. (d) Scoring of the predictors sorted in descending order. IM: image moment. AN: AlexNet feature. BoF: SURF features. *L*_1_|*L*_2_: ratio of long axis over short axis. < *I* >: average intensity. *P/A:* perimeter to area ratio. *σ_I_*: standard deviation of intensity. LF: long filamentous, SF: short filamentous, LS: large spherical, SS: small spherical, RD: rod-shape, UK: unknown.

The features were selected based on the following criteria: basic structural features of the particles (e.g. area, eccentricity) and Hu’s moments were chosen because they are rotationally and translationally symmetric. For the features from AlexNet and SURF a feature selection approach was designed based on maximising the standard deviation across the different structural classes. This approach constitutes a more rational choice compared to simple principal component analysis (PCA), which does not typically take the information regarding the classes into account, therefore our method selects for predictors that have high potential for class discrimination. This data reduction narrowed down the number of predictors to 6 for AlexNet and 6 for SURF. A total of 24 descriptors was finally chosen: 7 based on basic shapes (area, ratio of axis lengths, eccentricity, solidity, perimeter-to-area, mean intensity, standard deviation of pixel intensities), 5 of Hu’s image moments (Hu1, Hu4, Hu5, Hu6 and Phi4), 6 features obtained from the pre-trained convolutional neural network (CNN) AlexNet and 6 from a SURF bag of features. Figure 3(a) highlights the set of features used to train the random forest algorithm.

Panels described in Figure 3(b) indicate the spread of pairs of descriptors highlighting that some descriptors support classification across specific classes better than other combinations. Here, the training dataset was used to build a scatter plot of pairs of descriptors with the knowledge of their true classification (see colour scheme). For instance, the pair of descriptors ***L_i_*/*L_2_*** and Area show a good separation between long filamentous (dark blue labels) and unknown structures (black labels). The confusion matrix (Figure 3(c)) highlights the effective true positive rate (TPR) and positive predictive values (PPV) across the different classes with an overall predictive power of 88.4%. We note that some long filamentous viruses are misclassified as small filamentous, some small filamentous are misclassifed as small spherical and that, on occasion, some unknown structures populate the predicted long and large spherical structure classes. Considering the simple shapes of these viruses, it is expected that a small fraction of particles are misclassifed as structures with close resemblance.

The scoring of the predictors presented in Figure 3(d) indicates the average predictive power of each individual predictor. A high score indicates a high capacity to discriminate between different classes. The scoring was performed by measuring the predictive power of the classification for many combinations of predictors and distributing the predictive power score across the predictors tested (see Materials and Methods for details). In other words, if a combination of 2 descriptors alone give a predictive power of 60%, a score of 30% is awarded to both individual predictors. This method was repeated and scores represent averages across > 13,000 different combinations of descriptors.

### Structural details of an NDV virus population

We analysed a total of ~6,500 particles using MiLeSIM and established that 49.7% of NDV particles presented a filamentous shape whereas the large spherical, small spherical and rods represent 18.6%, 7.8% and 7.3% of the total population, respectively (Figure 4(a)). In addition to structural classification, the high-resolution images also permitted a dimensional analysis to be performed at the single particle level. We applied spherical ELM analysis to the large spherical viruses to extract the particle radius; backbone extraction to the short and long filamentous particles, to estimate the particle length; measured the equivalent radius of the particles for small spherical; and designed a model fitting for the rod structures. Figure 4 shows the distribution of structural parameters for each class. We observe that both long filamentous and large spherical are well described by a Gamma distribution whereas the small filamentous and small spherical are well described by a Gaussian distribution. The model-fitting applied to the rod-shaped particles (see materials and methods and ***Figure Supplement 2*** for details) allows the extraction of both the width and length of each particle. Therefore, it is possible to plot the distribution of structural parameters as a contour plot (Figure 4(c)).

**Figure 4.**
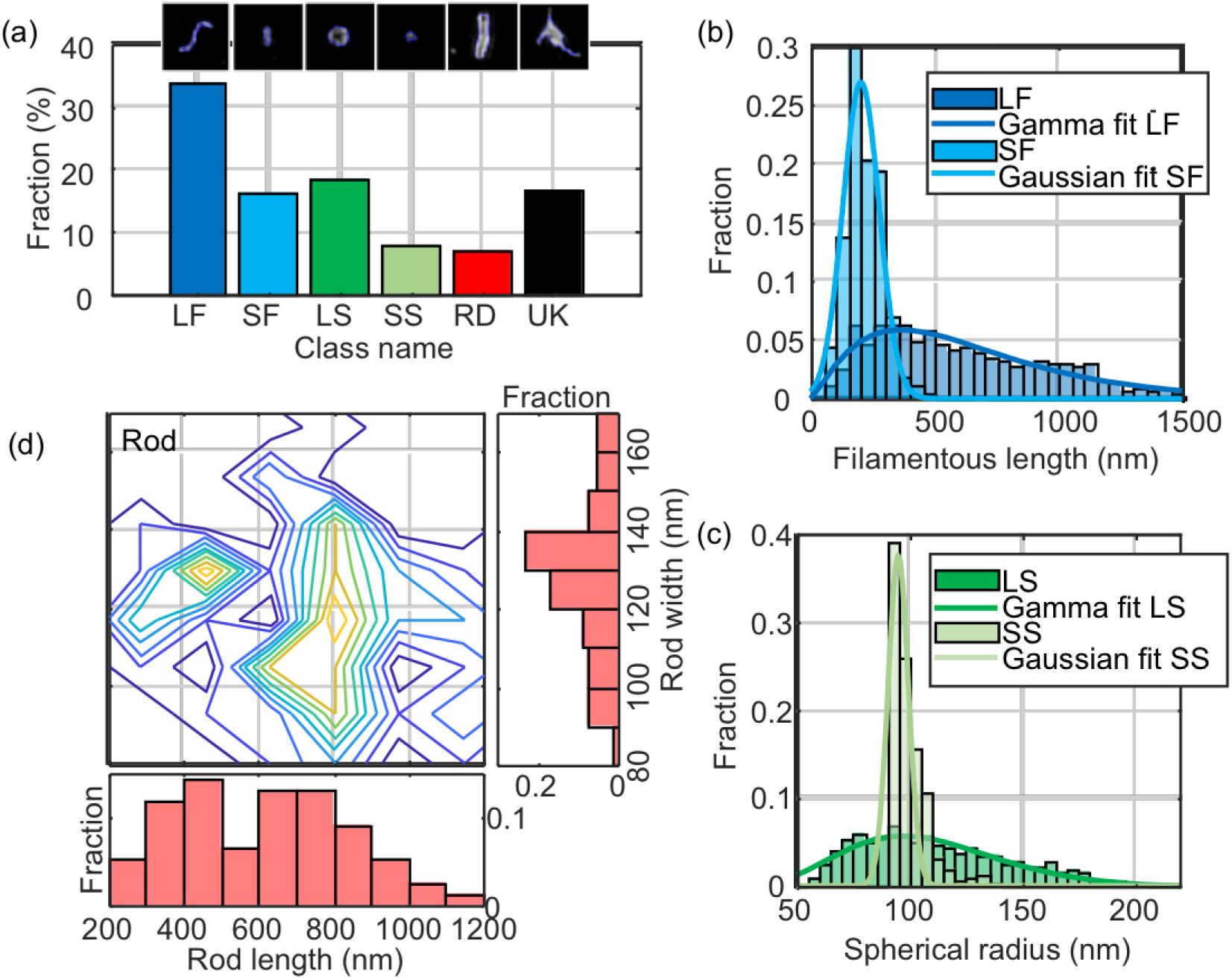
Quantitative analysis of NDV. The distribution of structural parameters for all classes was obtained from a total of ~6,500 virus particles. LF: long filamentous, SF: short filamentous, LS: large spherical, SS: small spherical, RD: rod-shape, UK: unknown.

We estimated the mean and standard deviation of the structural parameters from the distributions and obtained: <*L_LF_*> = *650* ± *430 nm*, <*L_SF_*> = *200* ± *100 nm*, <*D_LS_*> = *210* ± *70 nm*, <*D_SS_*> = *190* ± *10 nm*. For the rod-shaped particles, we observed that the width <*W_RD_*> = *135* ± *30 nm* and the length <*L_RD_*> = *610* ± *350 nm* (all rounded to two significant figures, ± represents the standard deviation of the distribution). These values are distributed around two populations as shown on the contour plot in Figure 4(d). It may first appear surprising that the distribution of radius of small spherical lies within that of the large spherical but in the ELM analysis the broadening of the image structure due to the finite optical resolution is effectively removed by taking into account the point spread function (PSF). It should be noted that the small spherical distribution is centred on the value of optical resolution of our SIM microscope, which indicates that the small spherical structures can be considered as point-like structures.

### MiLeSIM is capable of assaying influenza strains used for vaccine production in purified and non-purified samples from the production line

We performed the approach on a range of Live Attenuated Influenza Virus (LAIV) samples and classified the particles using the same classifier as for NDV. The LAIV virus population was dominated by spherical structures (> 60%). Figure 5 shows the distribution of particle sizes for four virus strains: a B-Victoria subtype (B/Brisbane/60/2008), a B-Yamagata subtype (B/Phuket/3073/2013) and two subtype A H1N1 strains (A/South Dakota /06/07 and A/Bolivia/559/2013). The fractions of small and large spherical particles are shown, as well as the equivalent radii and representative images of the viruses. It is clearly seen that B-Victoria particles consist of mostly large hollow particles with an equivalent radius of ~130 nm, a value that is in good agreement with the ELM analysis. Overall this leads to a particle radius of ~100 nm and a resolution of ~90 nm (***Figure Supplement 1***). In contrast, the B-Yamagata strain shows small and large particles of equal amount, indicating that the particles sizes are distributed around the region of overlap between small and large particles. This is confirmed by the nearly identical equivalent radius distributions.

**Figure 5.**
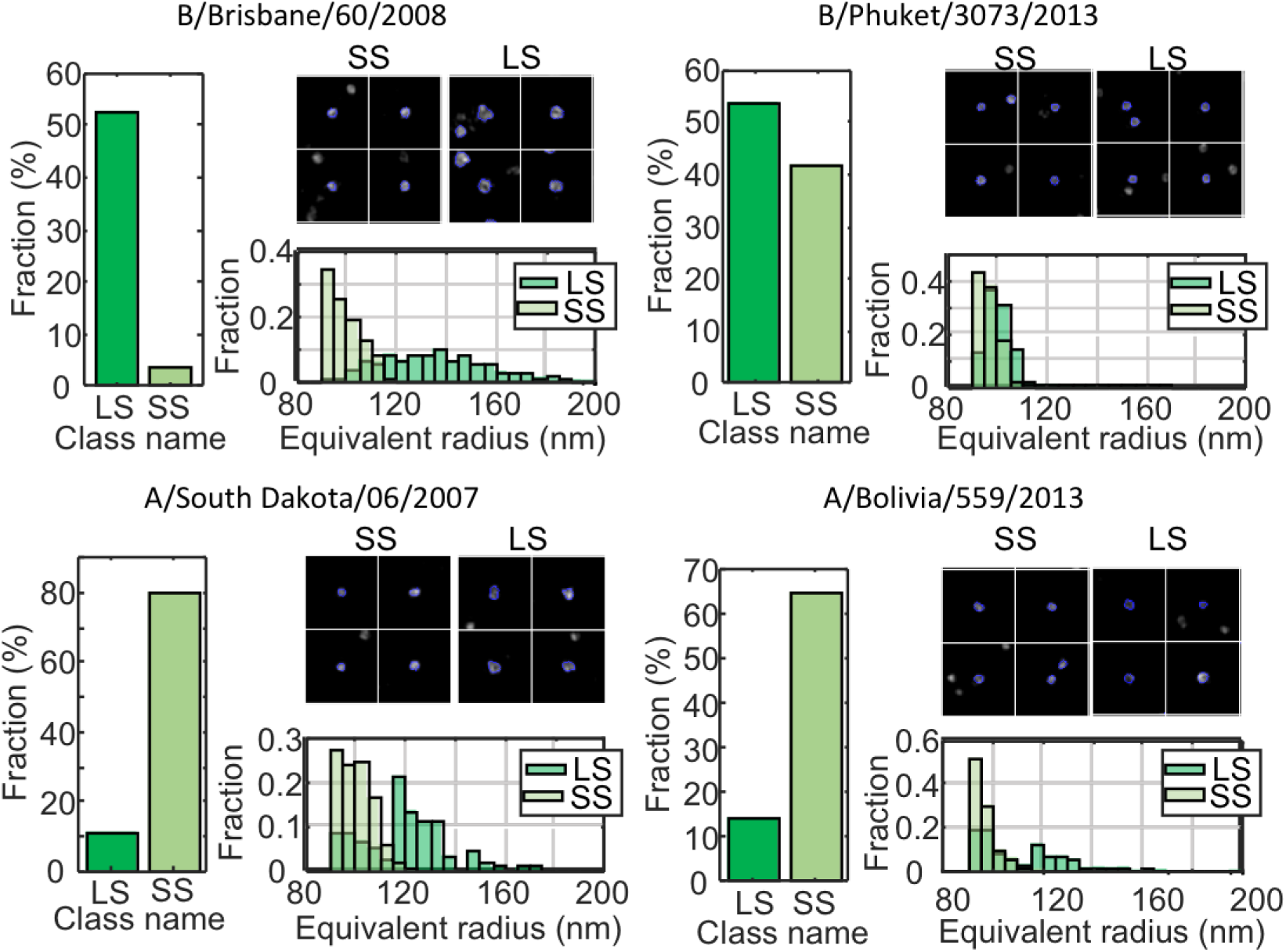
MiLeSIM approach applied to Live Attenuated Influenza Virus (LAIV). 2 types of B and A viruses were analysed here. The population was dominated by small and large spherical particles. The distributions of equivalent radius are shown here for both the large and small spherical for direct comparisons. The number of particles analysed were N=3,821, 4704,1,062 and 1,756 for B/Brisbane/60/2008, B/Phuket/3073/2013, A/South Dakota/06/2007 and A/Bolivia/559/2013 respectively.

Both A strains appeared clearly dominated by small spherical particles with sizes close to the resolution limit of our imaging. However, our high-throughput approach reveals subtle differences in the distribution of small spherical structures where the A/South Dakota viruses appear more heterogeneous (standard deviation ~10 nm), whereas the A/Bolivia viruses are sharply distributed (standard deviation ~4 nm).

We also investigated the potential of directly imaging pool harvested fluid (PHF). LAIV are commonly propagated in embryonated hens’ eggs where progeny viruses are released into the allantoic fluid of the egg. This fluid is harvested from numerous eggs and pooled. This constitutes a very basic and commonly used virus material. It is easy to produce and does not undergo any downstream purification. Consequently, PHF is impure, containing a variety of egg-derived impurities. The high molecular specificity of fluorescence microscopy allowed us to visualize the structure of the viruses with the same image quality directly in PHF despite the presence of a large amount of impurities (***Figure Supplement 2***). MiLeSIM therefore enables the study of unpurified samples and allows probing the virus production at any intermediate levels of production and purification. This constitutes a strong advantage over electron microscopy (EM) techniques which require the use of highly purified samples and elaboration preparation protocols.

## Discussion

We have demonstrated the potential of high-throughput imaging of virus structures, taking advantage of the optimal combination of speed and resolution afforded by the TIRF-SIM imaging method. TIRF-SIM provided sufficient resolution to identify, discriminate and analyse individual viral structural classes with high specificity, even in non-purified samples. Our approach combines machine learning to classify NDV viruses, followed by a model-based or direct quantification of virus structural parameters. The method yielded similar results both in purified samples and in samples from unfiltered PHF offering promise for use as an assay during virus production. We were able to image up to 500 particles / second at 90 nm resolution, vastly increasing throughput to alternative super-resolution methods or EM, which is traditionally used to analyse virus structure. Furthermore EM does not feature the specificity to analyse virus samples in their aqueous, unaltered forms. We were able to observe large structural variabilities in the NDV population and also between different strains of LAIV.

Our particular classification uses random forest with a selection of descriptors from simple shape parameters, rotational and translational invariant image moments and features from AlexNet and common feature for image recognition such as SURF. The overall predictive power of the approach is ~88.4% and the mis-classifcation occur between classes that are similar (between small spherical and small filamentous for instance). The structural parameters that we extract from the model fitting are precise beyond the image resolution as they take into account the finite optical resolution. This therefore reveals subtle differences in populations such as the two sub-classes observed in the rod-shaped class. This approach will be beneficial especially when heterogeneous populations are present and need to be quantified. In future, such information can be correlated with functional characteristics of produced virus classes and production parameters can accordingly be optimised. The approach thus holds great promise for the production of virus-based therapeutics. We note, however, that the methods presented are generally applicable to other systems and they are not restricted to a particular type of fluorescence microscopy, SRM or not.

## Materials and methods

### Sample preparation

The purified NDV samples were prepared on cover slips as previously described (Laine et al., 2015). Briefly, viruses were adhered on poly-L-lysine-coated Ibidi 8-well dishes, fixed, permeabilised and immuno-labelled with primary antibodies (mouse anti-hemagglutinin-neuraminidase HN, Abcam, UK) followed by secondary labelling (goat anti-mouse labelled with Alexa Fluor 647 for *d*STORM, with Alexa Fluor 488 for TIRF-SIM and with ATTO647-N for STED, Abcam, UK). The LAIV samples were prepared identically but using F16 mouse antibody for B-Victoria, Infa0121 mouse antibody for B-Yamagata and FY1 human antibody for A South Dakota and A Bolivia. The corresponding secondary antibodies were used (donkey anti-mouse DyLight 488 labelled or rabbit anti-human DyLight 488 labelled antibodies, ThermoFisher). All virus samples originated from the master virus bank (MVB) and are therefore highly purified, unless indicated in the text, where the direct pool harvest fluid (PHF) was used.

### TIRF-SIM, STED and *d*STORM imaging

Our custom-built TIRF-SIM system was described previously (Young et al., 2016). We used an Olympus UAPON 100x TIRF NA=1.49 and an Orca Flash 4.0 camera, with a sample pixel size of 64 nm. A total of 9 SIM images were acquired (3 phases, 3 orientations) with a camera exposure time of 200 ms and ~250 *μ*W of 488 nm laser, measured at the back aperture of the objective. The SIM images were obtained using the reconstruction code provided by Dr Lin Shao (Shao et al., 2011), providing images with doubled resolution and 32 nm final pixel size using a Wiener filter of 0.01. The STED imaging was performed on our custom-built STED microscope as described previously (Mahou et al., 2015; Curry et al., 2017). The *d*STORM imaging was performed on a custom-built single-molecule microscope previously described (Ströhl et al., 2017; Wong et al., 2017) and with and mercaptoethylamine (MEA) buffer as previously described (Laine et al., 2012). The *d*STORM image reconstruction was carried out using rapi*d*STORM 3 (Wolter et al., 2012).

The resolution achieved by the TIRF-SIM microscope was assessed by identifying the edge of the spatial frequency support using the SIMcheck plugin (Ball et al., 2015), as shown in ***Figure Supplement 1***. For STED microscopy, the resolution was estimated from cross-sections of 20 nm beads and reporting the full width at half maximum (FWHM). The *d*STORM resolution reported here was obtained from the FWHM of the localization precision, estimated by Thompson et al. (2002).

### Classification

All segmentations, features extractions and classifications were performed using MATLAB (Mathworks). A general diagram of the method is shown in ***Figure Supplement 1***. The segmentation was obtained by an initial Otsu binarization and refined by active contour. This allowed a better outline of the particles and efficient separation of particles in close proximity. The particles that were judged too small or too dim to be particles were excluded from further analysis. The basic shape features were extracted using the function regionprops. Hu’s image moments were computed from the 71×71 pixels particle image centred on the centre of mass of the particle. The absolute values of the logarithm of the moments were used in the classification. For the features obtained from AlexNet (Krizhevsky et al., 2012), the individual 71×71 pixels images were resized to 227×227 pixels and used as all three color layers of the RGB images taken by AlexNet. Then, feature extraction was performed using AlexNet as a pre-trained network. 4096 features were obtained and data reduction was performed to limit the number of predictors used. For this, the features were averaged within each individual class and the standard deviation of every feature across the classes was computed. The 6 features with the highest standard deviation was selected. For the SURF features, first a bag of visual words was created from the training dataset, this bag was then used to check the presence of visual words in the 71×71 pixels images of individual particles. Similarly to AlexNet features, we selected only the 6 visual word features with the highest standard deviation across the different classes for classification. This allowed the computation of a total of 24 features for ML.

The classification was performed using a random forest algorithm. The training dataset was made of 370 manually labelled individual particles and was used to train the random forest across 60 epochs. The classification was validated by 10-fold cross validation on the same dataset. The confusion matrix obtained from this cross-validation is show in Figure 3. At the training stage, the training dataset was augmented 5-fold by transforming the images with image translation and rotation randomly picked between 0 and 1 pixels and between 0 and 360 degrees respectively.

### Feature scoring

The features were scored by computing the predictive power of the random forest trained on the training dataset but with only subsets of features. Out of the 24 features all combinations of 2, 3, 4, 24, 23 and 22 features were tested corresponding to a total of 13,227 combinations of features. For each combination of features, the predictive power obtained was split equally across the different features used, producing a "local" predictive power for each feature. This local score was average across all combinations using a specific feature to obtain the global score.

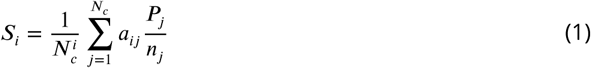

Where *S_i_* is the global score of the feature *i, 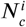* is the total number of combinations tested involving feature *i*, *a_ij_* is a factor reflecting the presence of the feature *i* in the combination *j*. *a_ij_* is equal to 1 if i is present in *j*, 0 otherwise. *P_j_* is the predictive power of the combination *j*, *N_c_* is the total number of combination tested and *n_j_* is the number of features present in the combination *j*.

### Quantitative analysis

All quantitative analyses were performed using MATLAB (Mathworks). The length of the filamentous structures were extracted by measuring the geodesic distance along the skeletonized image of the filament. The ELM analysis is freely available (Manetsberger et al., 2015) and the code was adapted to insert within the workflow of our approach. The equivalent radius *r* of the small spherical particles were simply calculated from the area *A* of the segmented particle.

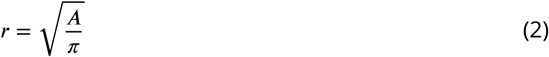

The image model for the rod-shaped particles is presented in ***Figure Supplement 1***. Briefly, the backbone of the particle was extracted by image thinning and then dilated by a disk-shaped kernel of radius equal to half of the width of the rod. The length of the rod could be adjusted by shortening the ends of the backbone or by extrapolating it outwards to lengthen it. The interior pixels of the image obtained were removed to leave the outline of the particle shape. This outline was then convolved with a Gaussian kernel in order to take into account the effect of the image resolution (here 90 nm). The intensity, the width and length of the model image were adjusted to minimize the sum of the square difference of intensity *χ*^2^.

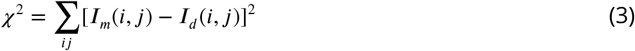

Where *i* and *j* refer to the indices in the image, *I_m_*(*i, j*) is the image model, and *I_d_*(*i, j*) is the original image.

## Acknowledgements

RFL and GG were supported by funding from MedImmune Ltd. We thank C. F. van der Walle for critical reading of the manuscript. CFK acknowledges funding from EPSRC, the Wellcome Trust, the UK Medical Research Council, MRC, MedImmune, and Infinitus (China), Ltd. RFL acknowledges a BBSRC TRDF grant (BB/P027431/1).

---

**Figure 1-Figure supplement 1.**
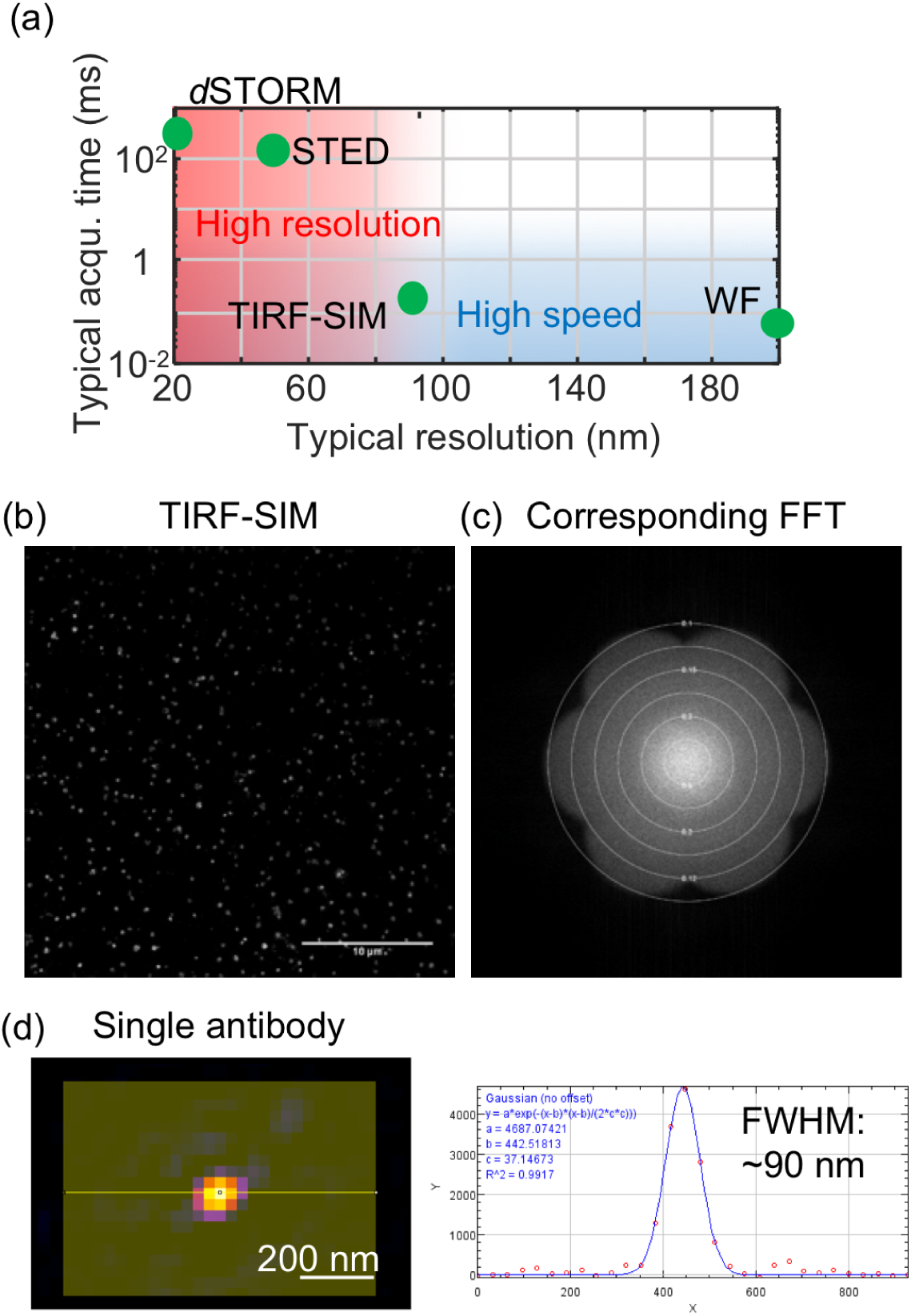
Resolution in SRM. (a) Typical spatial resolution and acquisition times for the imaging of NDV structures highlighting the trade-off between speed and resolution. (b) Representative TIRF-SIM image obtained from purified B-Victoria LAIV and its corresponding Fourier transform (c). The Fourier transform highlights the resolution ~90 nm. The plot was obtained using the SIMcheck plugin (Ball et al., 2015). (d) Image and cross section of a single secondary antibody labelled with DyLight 488, showing a FWHM of ~90 nm. FFT: Fast Fourier transform.

**Figure 2-Figure supplement 1.**
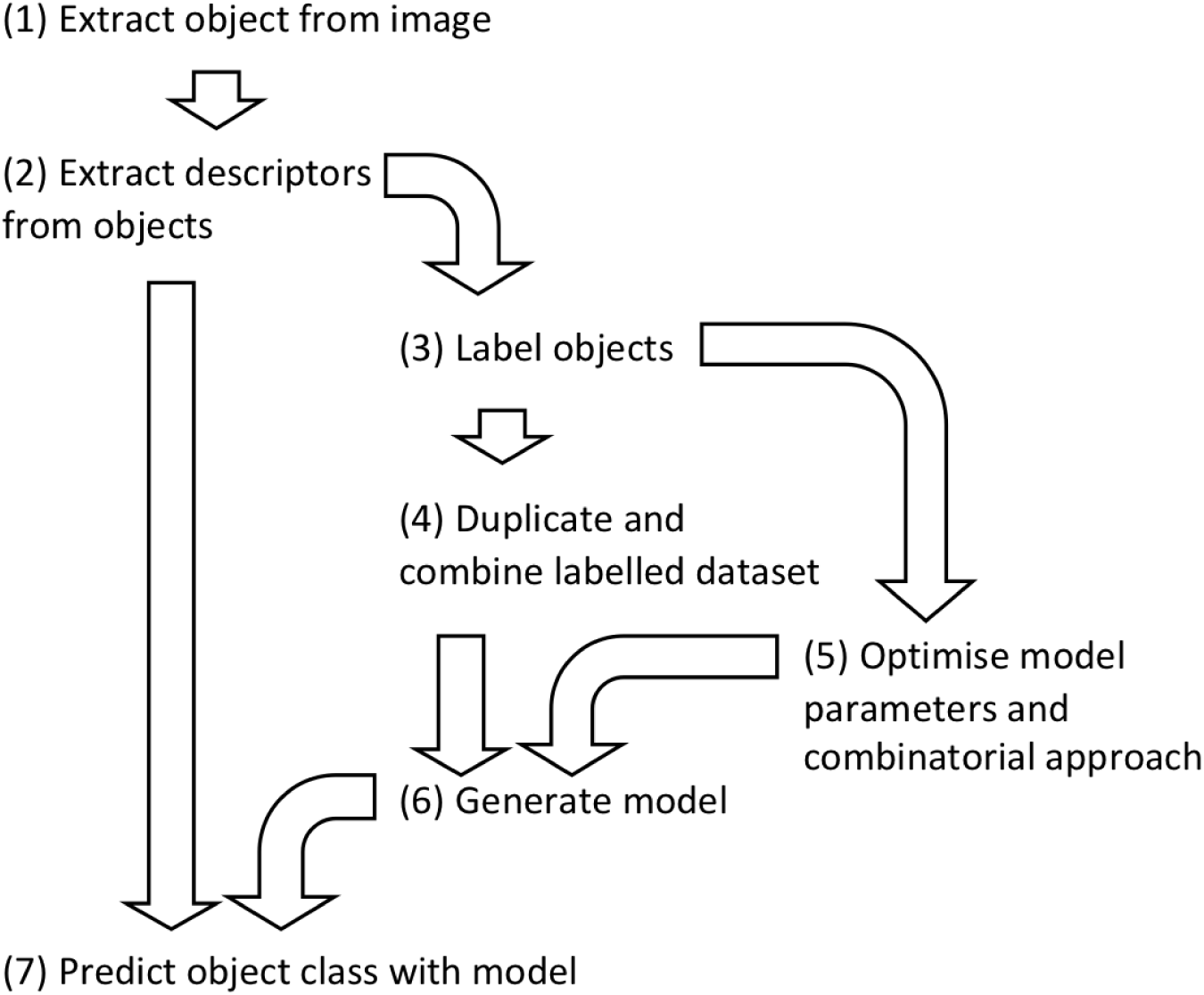
Flowchart describing the machine learning pipeline used here for image classification. A strong emphasis should be put on the choice of descriptors and the quality of the manual annotation (training dataset) prior to classification, as this will largely determine the quality of the classification.

**Figure 2-Figure supplement 2.**
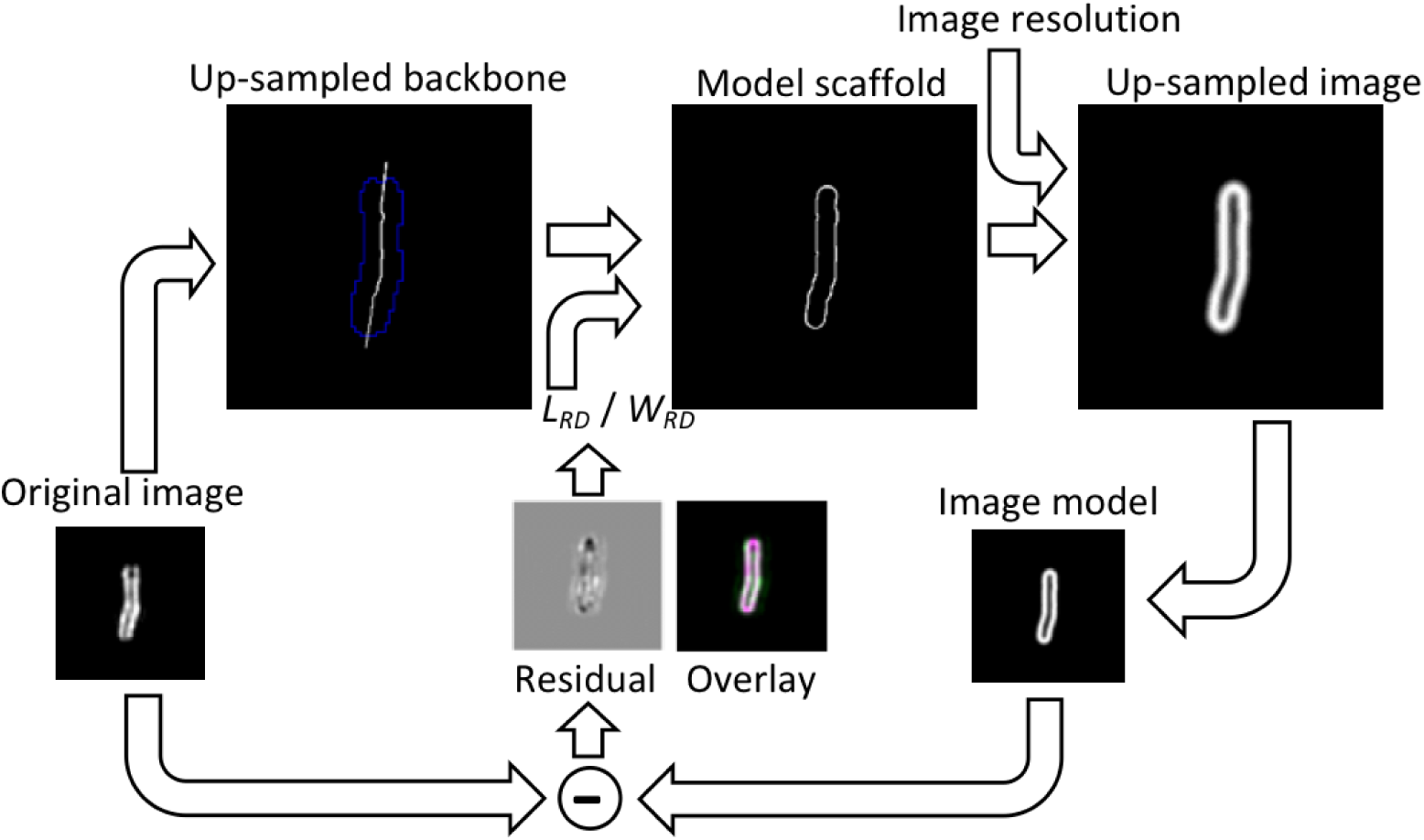
Image model for the analysis of the rod-shaped particles. The original image is segmented and thinned to obtain the backbone of the particle. The backbone is up-sampled and interpolated outside the particle. It is then used to compute the model scaffold. For this, each end of the backbone is independently grown or reduced to adjust its length, hence *L_RD_ = L_tot_ – L*_1_ − *L*_2_, where *L_tot_* is the maximum length of the extended backbone, and *L*_1_ and *L_2_* are the adjusted distances by which the backbone length is adjusted on each end respectively. Then, the image is dilated by a disk-shaped kernel of radius equal to half *W_RD_*. The outline of this image gives the model scaffold. The scaffold is then convolved with a Gaussian kernel representing the effect of image resolution (here 90 nm) and the image is down-sampled again to the original image size. The optimal *L_RD_* and *W_RD_* are those that minimize the difference image and the χ^2^.

**Figure 5-Figure supplement 1.**
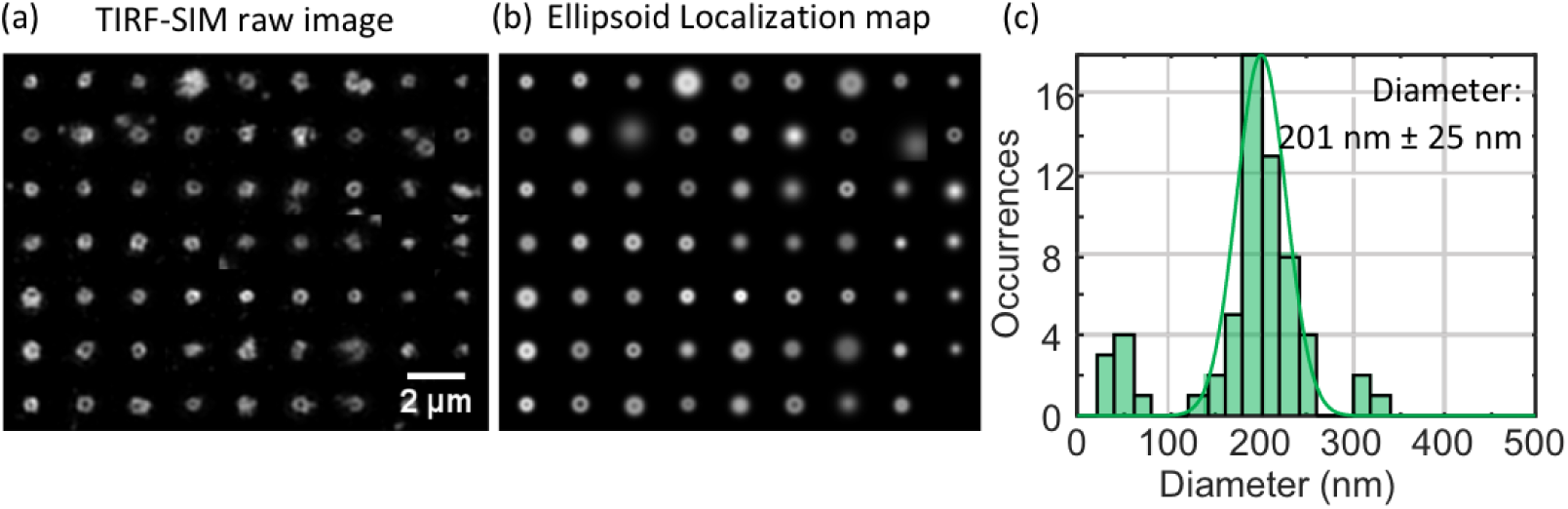
Spherical ELM analysis of the B-Victoria LAIV virus. Raw SIM images (a) and ellipsoid localization map (b). The distribution of diameters and a Gaussian fit are shown in (c). The diameter and error on the fit are the mean diameter and standard deviation obtained from the Gaussian fit.

**Figure 5-Figure supplement 2.**
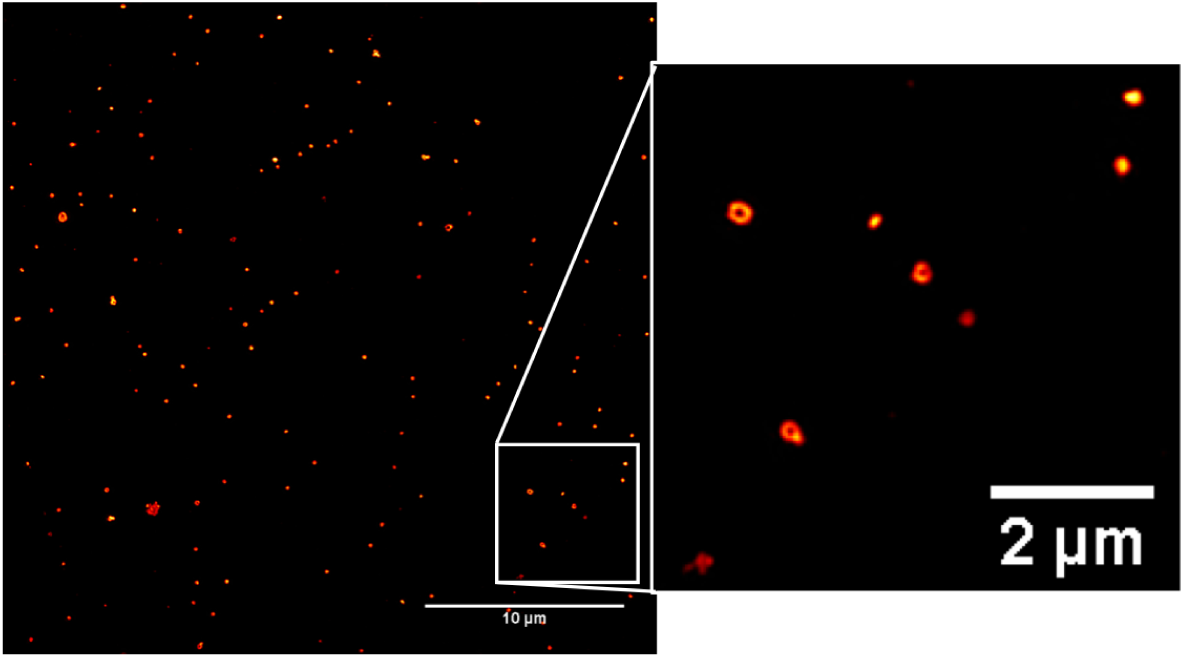
TIRF-SIM images obtained from pool harvested fluid (PHF). The images acquired here using PHF show an identical image quality as with highly purified samples.

